# Ecological selection of siderophore-producing microbial taxa in response to heavy metal contamination

**DOI:** 10.1101/163774

**Authors:** Elze Hesse, Siobhán O’Brien, Nicolas Tromas, Florian Bayer, Adela Lujan, Eleanor van Veen, Dave J. Hodgson, Angus Buckling

**Affiliations:** ESI & CEC, Biosciences, University of Exeter, Penryn Campus, Cornwall, TR10 9FE, UK.; Institut für Integrative Biologie, ETH Zürich, Universitätstrasse 16, Zürich, 8092, Switzerland.; Département de sciences biologiques, Université de Montréal, 90 Vincent-d’Indy, Montréal, QC H2V 2S9, Canada.; CIQUIBIC, Departamento de Química Biológica, Facultad de Ciencias Químicas, CONICET, Universidad Nacional de Córdoba, Córdoba, Argentina.; Camborne School of Mines, CEMPS, University of Exeter, Penryn Campus, Cornwall TR10 9FE,UK.; CEC, University of Exeter, Penryn Campus, Cornwall, TR10 9FE, UK.

**Keywords:** Adaptation, Detoxification, Ecological species sorting, Evolution, Metal tolerance, Public good dynamics, Remediation, Selection

## Abstract

Some microbial public goods can provide both individual and community-wide benefits, and are open to exploitation by non-producing species. One such example is the production of metal-detoxifying siderophores. Here, we investigate whether heavy metals select for increased siderophore production in natural microbial communities, or whether exploitation of this detoxifying effect reduces siderophore production. We show that the proportion of siderophore-producing taxa increases along a natural heavy metal gradient. A causal link between metal contamination and siderophore production was subsequently demonstrated in a microcosm experiment in compost, in which we observed changes in community composition towards taxa that produce relatively more siderophores following copper contamination. We confirmed the selective benefit of siderophores by showing that taxa producing large amount of siderophores suffered less growth inhibition in toxic copper. Our results suggest that ecological selection will favour siderophore-mediated decontamination, with important consequences for potential remediation strategies.

**Authorship:** EH, SOB, AL, DJH, EvV, AB conceived and designed the experiment. DJH provided new perspectives. EH, SOB, FB, AL collected the data. EH, FB, NT, DJH carried out the data analyses. EH & AB wrote the first draft of the manuscript, and all authors contributed substantially
to revisions.

**Data accessibility:** Upon acceptance, data presented in the manuscript will be made available on Dryad.

## INTRODUCTION

It is becoming increasingly apparent that many public goods benefit not only conspecifics but also other species. For example, many bacterial proteases show extracellular activity, providing potential nutritional benefits to neighbouring bacteria regardless of taxa (Suleman 2016); and immune-repressing molecules produced by parasitic nematodes provide a potential benefit to all co-infecting parasites (Maizels *et al.* 2001). Regardless of whether public goods are solely conspecific or also have interspecific benefits, there is the potential for non-producing cheats to outcompete producers assuming public good production carries some metabolic cost (Hamilton 1964; Hamilton & Axelrod 1981; Frank 1994). Hence, the evolution of costly public goods is crucially dependent on the extent to which benefits are reaped by the producer, other individuals carrying the public good gene and non-producers. While the evolution of public goods has been studied extensively within species, we know very little about how ecological species sorting acts to shape the production of interspecific public goods within natural communities. Here we combine surveys and experiments to determine how ecological selection acts on a microbial interspecific public good: siderophore-mediated heavy metal decontamination.

Heavy metals are ubiquitous components of the Earth’s crust, and large amounts have been released into the environment as a result of human activities (Nriagu & Pacyna 1988). Most heavy metals are toxic to microbes to varying degrees (e.g., Giller *et al.* 1998) and their presence can have a major impact on microbial communities (e.g., Gans *et al.* 2005). In the face of long-term selection imposed by heavy metals (Silver 1998), microbes have evolved mechanisms to cope with their toxicity, including metal reduction, reduced cell permeability and extracellular sequestration (Nies 1999; Bruins *et al.* 2000; Valls & De Lorenzo 2002). One such detoxification mechanism is the production of siderophores. While the canonical function of siderophores is to scavenge poorly soluble iron (Ratledge & Dover 2000), bacteria also use these secreted molecules to bind heavy metals (Braud *et al.* 2010; Schalk *et al.* 2011). Siderophore production can be induced by the presence of non-iron metals (Hofte *et al.* 1993; Teitzel *et al.* 2006), which they bind with various affinities (Braud *et al.* 2009). These siderophore-metal complexes, unlike siderophore-bound iron, are unable to enter bacterial cells, thereby reducing the concentration of free toxic metals in the environment (Schalk *et al.* 2011). This has led to the suggestion of adding siderophores or siderophore-producing microbes to remediate heavy metal contaminated environments (Diels *et al.* 1999; Gadd 2000; Rajkumar *et al.* 2010; O’Brien & Buckling 2015). However, to understand how siderophores may both contribute to the natural decontamination of environments and the long-term success of remediation strategies, it is crucial to determine how toxic metals affect selection, within and between species, for siderophore production in natural communities. Note that we do not simultaneously address within species selection alongside ecological selection, largely because the genetic resolution of the sequencing methods used to identify taxa is only at the level of genus.

Given their detoxifying effect, increasing metal toxicity might be expected to result in ecological species sorting in favour of species with greater siderophore production. However, the production of siderophores in the context of decontamination not only benefits the producer (or its close relatives), but potentially also neighbouring cells, both con- and hetero-specific, in the community. Siderophore production is often associated with a fitness cost, hence selection may favour cells that produce fewer siderophores, but still receive the same detoxifying benefits of siderophore production from neighbours (West *et al.* 2007; O’Brien *et al.* 2014). Limited diffusion of such public goods (Kummerli *et al.* 2009; Kummerli *et al.* 2014) and positive assortment of producing cells resulting from spatial structure (Hamilton 1964; West *et al.* 2007; Mitri & Foster 2013; Ghoul & Mitri 2016; Pande *et al.* 2016) may, however, limit the community-wide benefits of siderophores and prevent overexploitation by non-producing cells (Kummerli *et al.* 2009; Oliveira *et al.* 2014). The situation is further complicated by the iron-scavenging function of siderophore production, which is also open to exploitation within (Griffin *et al.* 2004; Buckling *et al.* 2007; Lujan *et al.* 2015) and between species (Barber & Elde 2015; Galet *et al.* 2015). It is therefore unclear if siderophore production will increase or decrease in natural communities as a function of metal toxicity, or whether it will result in stable coexistence of producing and non-producing taxa (Hibbing *et al.* 2010; Cordero *et al.* 2012; Morris *et al.* 2012; Morris 2015; Estrela *et al.* 2016).

To investigate how heavy metal contamination affects ecological selection for siderophore production, we first confirm that siderophores can act as an interspecific public good in an *in vitro* siderophore-addition experiment. We then conducted a simple survey along a natural gradient of metal contaminated soil. We correlated total metal concentration with species composition and estimates of siderophore production determined from the proportion of culturable bacteria that show detectable extracellular iron-chelation *in vitro*. We then conducted an experimental study in compost communities to determine causal links between metal contamination and siderophore production

## METHODS

### Siderophores as interspecific public goods

To test whether siderophores can act as interspecific public goods we quantified whether the presence of yersiniabactin, a copper-chelating siderophore produced by *Yersinia pestis* (Chaturvedi & Henderson 2014), can ameliorate growth of a non-producing strain of *Pseudomonas aeruginosa* in toxic copper broth. Both species are Gram-negative opportunistic pathogens belonging to the γ Proteobacteria, and share many, potentially cooperative, functional traits (e.g., type III secretion system; Rundell *et al.* 2016). We inoculated ~10^4^ colony forming units (CFU) of a producing strain of *P. aeruginosa* (PA01) and an isogenic non-producing mutant strain (PA01*ΔPvdDΔPchEF*) in isolation into four replicate micro-centrifuge tubes, containing 900 μl of KB broth with or without 0.6 mM CuSO_4_, which reduces relative non-producer fitness (O’Brien *et al.* 2014). In addition, ~10^4^ CFUs of either strain were inoculated in 900 μl KB supplemented with equal molarities of CuSO_4_ and yersiniabactin, which typically binds to Cu(II) at a 1:1 ratio (Koh & Henderson 2015). Copper is a common heavy metal (Nriagu & Pacyna 1988), and is one of dominant metals found at our field site (Fig. 2A); hence this is why we used copper sulphate in our *in vitro* assays. Bacterial cultures were horizontally shaken at 37°C for 24hr, after which serial-diluted culture was plated onto KB agar to obtain cell densities and calculate population growth: *m*=ln(N_f_/N_0_), where N_0_ and N_f_ are the initial and final bacterial densities.

**Figure 1.**
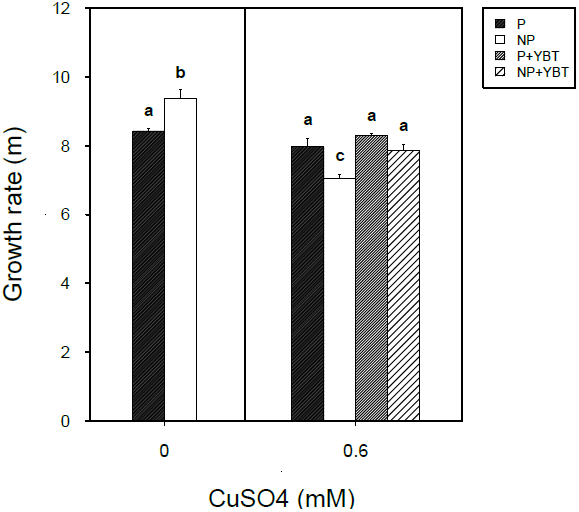
Siderophores act as an interspecific public good in toxic copper broth. Growth rate (m) of a non-producer (NP) and producer (P) strain of *Pseudomonas aeruginosa* in control (0 mM CuSO_4_) and copper-contaminated broth (0.6 mM CuSO_4_) with (YBT) and without yersiniabactin. Letters denote significant differences based on post-hoc Tukey contrasts.

**Figure 2.**
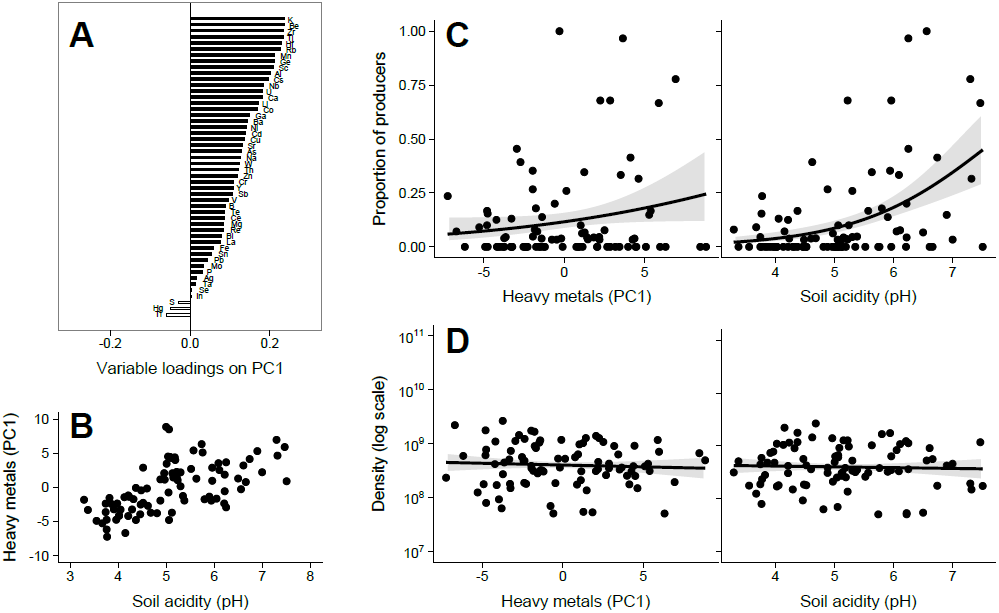
The effect of soil acidity and heavy metal contamination on microbial abundance and siderophore production in natural soils. **(A)** Heavy metal loadings onthe first principal component (PC1), which explained 27% of the observed environmental variation; **(B)** Positive correlation between soil acidity (pH) and heavy metal contamination (PC1); **(C)** Proportion of siderophore producers and **(D)** microbial density (log_10_-transformed bacterial cells g^-1^ soil) as a function of heavy metal contamination and soil acidity. Lines and shaded area depict the fitted relationships ± standard error.

To test whether non-producers can exploit the detoxifying benefit of yersiniabactin, we used a two-way ANOVA with strain (2 levels) x treatment (3 levels: KB-, copper- and yersiniabactin + copper broth) as explanatory variables.

### Natural microbial communities

#### Soil collection and characterization

Soil samples were collected in a former poly-metallic mining area situated in the Poldice Valley (N: 50º14.56; W: 5º10.10) in Cornwall (UK). The valley is rich in heavy metals, as apparent from the significant production of tin, copper, arsenic, lead, zinc and silver during the 18-19^th^ centuries (Burt 1998), which fuelled the industrial revolution. The area, once referred to ‘the richest square mile to be found anywhere on earth’, is no longer worked leaving a legacy of untreated mining waste. 94 samples were collected by pushing sterile bulb planters into the ground near calciners, chimneys, slag heaps and regenerated areas, representing a wide range of metal contamination. The upper part of the soil core was discarded to rule out possible contamination from the ground surface. Samples were then transferred to sterile 50 millilitre (ml) falcon tubes and stored at 4°C until further processing. Samples were sieved (1 millimetre mesh) prior to DNA extraction and quantification of pH and heavy metals.

Quantification of heavy metals and metalloids (e.g., Fe, Cd, Cr, Cu, Mn, Hg, Ni, Ti, V, Zn, Pb, Sn, As) was carried out by ALS global (Loughrea, Ireland), using an aqua regia digest (EPA 3050b). To assess the total content of these determinants, samples were analysed using emission spectroscopy (ICP-OES). For each sample, we quantified pH by suspending 1 gram (g) of soil in 5 ml of 0.01M CaCl_2_ (Hendershot & Lalande 2008). The suspension was shaken for 30 minutes (min) and left to stand for 1 hour (h), after which the pH of the supernatant was measured using a Jenway 3510 pH meter (Stone, UK).

#### Siderophore production

The relationship between siderophore production, soil acidity and metal contamination was tested by screening a subset of clones for siderophore production. Siderophore production was necessarily measured under common garden conditions, to avoid the confounding effect of environmental variation if conducted *in situ*, causing both differential induction of siderophore production as well as different metal-chelating activities of the different soil types themselves, which could directly affect the siderophore assay. For each sample, 1 g of soil was transferred to 6 ml of M9 solution in 30 ml glass vials, which were shaken for 2h at 28°C and 180 rpm, after which serial-diluted supernatant was plated onto LB agar. Thirty individual colonies per sample were randomly selected and grown for 48h independently in 200 microliter (μl) KB broth at 28°C. A 2 μl sample from each colony was then spotted on blue-tinted iron-limited CAS agar plates (Schwyn & Neilands 1987) using a pin replicator. Plates were incubated at 28°C for 48h, after which we scored the presence of orange halos, a qualitative indicator of siderophore secretion, to obtain an estimate of the proportion of siderophore-producing clones in each community.

#### DNA extractions and real time PCR

To determine how microbial abundance and community composition varied across soils we extracted genomic DNA from 250 milligram (mg) soil per sample, using the MoBio Powerlyzer PowerSoil© DNA isolation kit (Carlsbad, CA, USA), following the manufacturer’s protocol with the bead beating parameter set to 4500 rpm for 45 seconds (s). The integrity of DNA was confirmed using 1% TAE agarose gels stained with 1x Redsafe DNA Stain (20 000X): based on quality checks, 5 samples were discarded from subsequent analyses, yielding 89 DNA samples in total.

Community density was quantified using real-time PCR (StepOnePlus Real-Time PCR, Applied Biosystems, Foster City, CA, USA) on 1:10 and 1:100 diluted samples with primers 16S rRNA 338f (ACT CCT ACG GGA GGC AGC AG) and 518r (ATT ACC GCG GCT GCT GG) (Øvreås & Torsvik 1998). Triplicates of each sample were run along gDNA standards (5 x 10^2-6^16S rRNA genes of *Pseudomonas fluorescens*) and non-template controls. All assays were based on 15μL reactions, using 1x Brilliant III Ultra-Fast SYBR^®^ Green QPCR Master Mix (Agilent technologies, Santa Clara, CA, USA), 150nM 338f and 300nM 518r primers, 300nM ROX and 100ng/μL BSA. Thermal conditions were set to: 3 min at 95°C for initial denaturation, followed by 40 cycles of 5s at 95°C and 10s at 60°C (after which fluorescent data were collected), followed by a melting curve at 95°C for 15s, 60°C for 1 min ramping up to 95°C in steps of +0.3°C for 15s. Melting curves and confirmation of non-template controls was analysed using StepOne Software version 2.3 (Applied Biosystems). Baseline corrections, Cq values and efficiencies (1.89 ± 0.07 and 1.89 ± 0.08 for standards and samples) were determined using LinRegPCR version 2016.0 (Ruijter *et al.* 2009). 16S rRNA gene quantities were calculates using the one point calibration method (Brankatschk *et al.* 2012). Bacterial counts were estimated by dividing 16S rRNA gene quantities by a mean copy number of 4.7 (ribosomal RNA operon copy number database rrnDB; version 5.1; February 16, 2017) (Stoddard *et al.* 2014), corrected for variation in dry weight.

#### Statistical analyses

Because of strong collinearity among heavy metals, we carried out a principal component analysis (PCA) on centred and scaled data. Most of the quantified metals loaded positively on the first principal component (PC1; Fig. 2A). For this reason, we used PC1 as a proxy for total metal contamination in all subsequent analyses. To test how PC1 and pH affect siderophore production we used individual generalized linear models (GLMs) with a binomial response variable and a quasi-binomial error structure. The effect of these environmental variables on microbial density was tested using individual GLMs on log_10_-transformed data.

#### Sequencing, OTU picking and diversity analyses

Library preparation and sequencing was performed by the Center for Genomic Research (University of Liverpool, Supplementary Methods).

Base-calling and de-multiplexing of indexed reads was performed by CASAVA version 1.8.2 (Illumina, San Diego, CA, USA) to produce 89 samples from the 1st lane of sequence data, which were trimmed to remove Illumina adapter sequences using Cutadapt version 1.2.1 (Martin 2011). The 3’ end of reads matching the adapter sequence over >3 bp was trimmed off. Reads were further trimmed to remove low quality bases, using Sickle version 1.200 with a minimum window quality score of 20. After trimming, reads <10 bp were removed. If both reads from a pair passed this filter, each was included in the R1 (forward reads) or R2 (reverse reads) file. If only one of a read pair passed this filter, it was included in the R0 (unpaired reads) file.

Sequences were processed with the default parameters of the SmileTrain pipeline (https://github.com/almlab/SmileTrain/wiki/), including reads quality and chimera filtering, paired-end joining, de-replication and *de novo* distribution-based clustering using USEARCH (version 7.0.1090, http://www.drive5.com/usearch/) (Edgar 2019), Mothur (version 1.33.3) (Schloss *et al.* 2009), Biopython (version 2.7), dbOTUcaller algorithms (Preheim *et al.* 2013; https://github.com/spacocha/dbOTUcaller, version 2.0) and custom scripts. We generated an OTU table that was filtered to minimize false OTUs using the filter_otus_from_otu_table.py QIIME script (Caporaso *et al.* 2010; http://qiime.org/; version 1.8) by removing OTUs observed <10. We assigned taxonomy, post-clustering, using the 97% reference OTU collection of the GreenGenes database (http://greengenes.lbl.gov; release 13_8). Taxonomy information was added to the OTU table using biom add-metadata scripts (http://biom-format.org/). A total of 8 604 074 sequences were obtained, ranging from 39 253 to 192 455 reads per sample, with a median of 91 646. This dataset was clustered into 45 891 OTUs.

Diversity calculations were based on non-rarefied OTU tables. β-diversity was calculated using the Jensen-Shannon Divergence (JSD) metric (Fuglede & Topsoe 2004; Preheim *et al.* 2013), which is robust to sequencing depth variation. The R ‘phyloseq’ package (version 1.19.1) (McMurdie & Holmes 2013; https://joey711.github.io/phyloseq/) was used to transform the OTU table into relative abundances, which were square-root-transformed into Euclidean metrics (Legendre & Gallagher 2001). Finally, we used Nonmetric Multidimensional Scaling (NMDS) plots (Shepard 1962; Kruskal 1964) to order bacterial community composition. Differences in community structure were tested using PERMANOVA (Anderson 2001), implemented using *adonis*() from the R ‘vegan’ package (version 2.4-1) with 999 permutations.

To confirm that pH and PC1 shape community structure, we used K-means partitioning algorithms (MacQueen 1967) implemented with *cascadaKM*() from the ‘vegan’ package with 999 permutations. K-means is a completely independent way of binning samples. We Hellinger-transformed (Rao 1995) the OTUs table using *decostand*(x. method=”hellinger”) and tested whether our samples naturally clustered into 2-10 groups based on their composition using the Calinski-Harabasz index (Caliński & Harabasz 1974).

To investigate how environmental variables contributed towards explaining variation in community composition, we used a multivariate regression tree analysis (MRT; Breiman *et al.* 1984; De’Ath 2002) for pH and PC1 separately, using *mvpart*() and *rpart.pca*() from the R ‘mvpart’ package (De’Ath 2007; Therneau *et al.* 2015). The OTU table was first Hellinger-transformed (Rao 1995) before carrying out the analyses (Ouellette *et al.* 2012). After 200 cross-validations (Breiman *et al.* 1984), we plotted and pruned the tree using the 1-SE rule (Legendre & Legendre 2012) to select the least complex model. We used *rpart.pca*() from the ‘mvpart’ package to plot a PCA of the MRT.

α-diversity was estimated using Shannon (Oksanen *et al.* 2010; ‘vegan’ package version 2.4-1) and Chao1 (Vavrek & Larsson 2010; ‘fossil’ package version 0.3.7) indices. We used *resample_estimate*() from the R ‘breakaway’ package (Willis & Bungle 2014, version 3.0) to account for sample size variability, setting the number of bootstraps to 500 with replacement. The relationship between α-diversity and environmental variables was tested using *betta*() from the ‘breakaway’ package, which accounts for statistical errors associated with estimating Shannon and Chao1 indices.

### Copper-addition experiment

#### Experimental design

To infer a causal relationship between toxic metals and siderophore production, we set up experimental communities in 90 millimetre Petri dishes containing 30g of twice-autoclaved 50% peat-free compost (Verve John Innes No. 1). Before sterilization, the natural microbial community was isolated by adding 40 g of fresh compost to 200 ml of M9 solution and incubating at 150 rpm at 28°C for 24h.

We established communities by inoculating twelve soil microcosms with 2 ml of soil wash (~2.4 × 10^7^ CFUs ml^-1^). Microcosms were placed in an environmental chamber at 26°C and 75% humidity for 24h, after which we supplemented six microcosms with 2 ml of filter-sterilised 0.25M CuSO_4_ or ddH_2_0. This concentration of CuSO_4_ hindered bacterial growth. Microcosms were then returned to the environmental chamber for a total of 6 weeks. After three weeks, another 2 ml dose of CuSO_4_ or ddH_2_O was added where appropriate. Samples of the community were taken prior to copper amendment and 3-6 weeks post-inoculation by transferring 1 g soil to 6 ml of M9 solution in 30 ml glass vials. Vials were shaken for 2h at 28°C at 180 rpm, after which soil wash supernatants were frozen at −80° C in 25% glycerol.

#### Siderophore and copper resistance assays

To quantify siderophore production, 24 individual clones per treatment-time combination were isolated by incubating serial-diluted soil wash on LB plates at 28°C for 48h. Individual colonies were then transferred to 2 ml of KB broth and grown for 48h at 28°C, after which the supernatant was assayed for the extent of iron chelation. Siderophore production was quantified using the liquid CAS assay described by Schwyn and Neilands (1987), with the modification that one volume of ddH_2_0 was added to the assay solution (Harrison & Buckling 2005). We used the following quantitative measure to obtain an estimate of siderophore production per clone: [1 − (A*_i_*/A*_ref_*)]/[OD*_i_*)], where OD*_i_* = optical density at 600 nanometre (nm) and A_i_ = absorbance at 630 nm of the assay mixture *i* or reference mixture (KB+CAS; A*_ref_*). Note that CAS assays performed in iron-limited KB (supplemented with 20 mM NaHCO_3_ and 100 μg ml^-1^ human apotransferrin) provided qualitatively similar results (data not shown).

All final time-point clones were grown at 28°C for 24h, after which ~10^4^ CFUs were inoculated into 96-well plate wells containing 200 μl of KB broth supplemented with or without a toxic dose of CuSO_4_ (6.17 mM). Clones were incubated statically at 28°C for 48h, and their OD was measured at 600 nm every 8-12h to quantify growth (Varioskan Flash plate reader, Thermo Scientific, Waltham, MA, USA).

#### Sanger sequencing of 16S rRNA of evolved clones

The 16S rRNA gene of all assayed final-time point clones was sequenced to confirm genus-level identity. In short, PCRs were performed in 25μL reactions containing 1x DreamTaq Green PCR Master Mix (2X) (Thermo Scientific), 200 nM of the 27F and 1492R primers and 3 μL of 1:100 diluted culture that had undergone 3 freeze-thaw cycles. The thermal cycling parameters were set to 94°C for 4 min, followed by 35 cycles of 1 min at 94°C, 30s at 48°C and 2 min at 72°C, and a final extension of 8 min at 72°C. Following Exo-AP clean-up, high quality samples were Sanger sequenced using the 27F primer (Core Genomic Facility, University of Sheffield).

The quality of all sequences was assessed using *plotQualityProfile*() from the R ‘dada2’ package (Callahan *et al.* 2016; version 1.3.0). Based on the obtained plots, sequences were trimmed in Genious (version 6.1.8) to achieve an overall quality score >35, by removing >20bp from the 5’ end and trimming the 3’ end to a maximum length of 700bp. Using Mother, sequences longer then 300bp were aligned to the Silva.Bacteria.Fasta database, and taxonomy was classified using the RDP trainset 14 032015 as a reference database.

#### Statistical analyses

The effect of copper on temporal changes in mean per capita siderophore production was tested using a linear mixed effect (LME; ‘lme4’ R package; Bates *et al.* 2014) model with copper x evolutionary time (3-6 weeks post inoculation) as fixed categorical effects and random intercepts fitted for each community (*n* = 12), and individual clones nested within communities (*n* = 24), to account for temporal dependencies.

We used NMDS ordination plots to depict pair-wise Bray-Curtis dissimilarities in genus-level composition between microcosms. To test whether treatments differed significantly in their composition we used PERMANOVA with 999 permutations, and tested for equality of between-treatment variance using permutation tests for homogeneity of multivariate dispersion.

To test for the effect of copper on metal tolerance, we used a LME model with *ln*(OD_Cu_/OD_KB_) as response variable, copper background as fixed effect and a random slope fitted for mean-centred hours: random=~(Hours)|Community/Clone. The model thus accounts for intrinsic differences between communities, and nested clones, in their ability to tolerate toxic levels of copper over time, and explicitly tests whether pre-adaptation to copper increases mean copper tolerance. To test whether tolerance was directly mediated by variation in siderophore production, we replaced ‘copper background’ with clone-specific siderophore production.

In general, full models were simplified by sequentially eliminating non-significant terms (*P* > 0.05) following a stepwise deletion procedure, after which the significance of the explanatory variables was established using likelihood ratio tests. In case of significant differences, Tukey contrasts were computed using the ‘multcomp’ package (Hothorn *et al.* 2008), with α < 0.05. We used *R* Version 3.1.3 for all analyses (R Development Core Team; http://www.r-project.org).

## RESULTS

### Foreign siderophores restore non-producers fitness in toxic copper medium

Strains responded differentially to copper (strain x treatment interaction: *F*_*2,16*_ = 18.06, *P* < 0.001), with non-producers outperforming producers in KB broth and *vice versa* in copper broth (Fig. 1). Crucially, the addition of yersiniabactin restored the growth of non-producers to levels comparable to that of siderophore-producers when propagated in toxic copper (Fig. 1).

### Microbial diversity, abundance and siderophore production along a natural heavy metal gradient

We found that the proportion of siderophore-producing isolates was significantly greater in more heavily contaminated soils (PC1: χ^2^ = 4.42; d.f. = 1, *P* = 0.04 Fig. 2C). Because contamination co-varied with soil acidity (Pearson’s product-moment correlation: *r* = 0.61, d.f. = 86 and *P* < 0.001; Fig. 2B), siderophore production also increased as a function of pH (χ^2^ = 28.16; d.f. = 1, *P* < 0.001; Fig. 2C). Neither pH nor PC1 significantly affected microbial abundance (GLM: *F*_*1, 87*_ = 0.01, *P* = 0.99 for PC1 and pH; Fig. 2D). Both environmental variables predicted community structure: samples with similar range values of pH (PERMANOVA: R^2^ = 0.087, *P* < 0.001) or PC1 (R^2^ = 0.065, *P* < 0.001) had similar community composition. Because the explanatory power of these variables was relatively low (Fig. S1 in Supplementary Information), we performed a K-means analysis, which showed that samples were naturally divided into 2 or 3 groups differing significantly in their PC1 or pH, respectively (Fig. S2 in Supplementary Information). We used MRT to confirm these findings and observed that R^2^ was highest when pH was used as explanatory variable (pH: R^2^ = 0.183 and PC1: R^2^ = 0.085; Fig. 3). Alpha diversity was largely independent of PC1, but varied as a function of pH (Fig. S3 in Supplementary Information; *P* < 0.001 for both indices).

**Figure 3.**
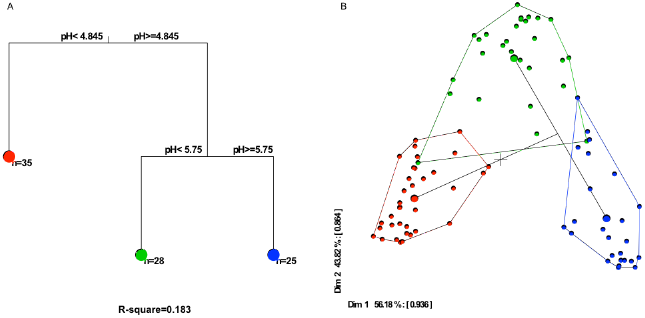
Community composition variation changes as a function of soil acidity. Multivariate regression tree (MRT) analysis was used to estimate the impact of soil acidity (pH) and heavy metals (PC1) on community structure, indicating that pH is the main environmental driver explaining variation in community structure. The most parsimonious tree (**A**) shows that the community could be divided into 3 different leaves (colored symbols) based on microbial abundance and composition. The composition within leaves is represented in a PCA plot (**B**), where small points represent individual samples and large points represent the group mean (within leaf). The most important taxa in each leaf are summarized in Supplementary Table S1.

### The effect of copper on siderophore production in experimental communities

Our assay of siderophore production along a natural gradient showed that siderophore production was greater in more contaminated soils. However, it remains unclear whether metals are a significant driver explaining variation in siderophore production. Notably, pH is an important predictor of soil bacterial diversity and composition (e.g., Fierer & Jackson 2006; Griffiths *et al.* 2011), and correlated positively with metal contamination, making any interpretations ambiguous. To determine a causal link between heavy metals and siderophore production, we carried out an experiment and characterised and measured siderophore production of multiple clones as well as their metal tolerance. We found that mean siderophore production was significantly greater in communities subjected to copper contamination (LME: copper effect: χ^2^=6.91; d.f. = 1; *P* < 0.01; Fig. 4A). Note, however, that overall siderophore production decreased through time (time effect: χ^2^= 16.02; d.f. = 1; *P* < 0.001) independent of treatment (time x treatment effect: χ^2^= 0.001; d.f. = 1; *P* = 0.98). Soil acidity marginally increased following copper contamination (mean pH ± SE after 3 and 6 weeks of incubation in control = 7.13 ± 0.05, 7.09 ± 0.02 and in copper = 6.90 ± 0.04, 6.60 ± 0.05), indicating that siderophore production was greater in more acidic compost.

**Figure 4.**
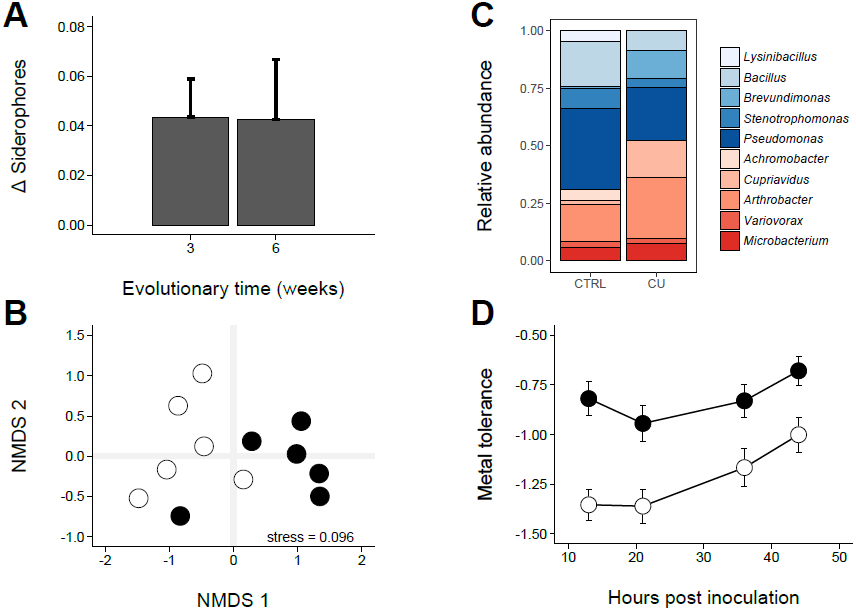
The effect of copper contamination on experimental microbial communities in compost. **(A)** Copper increases mean per capita siderophore production. Bars depict the mean difference in siderophore production ± 1 S.E. between copper-and control treatments after 3-6 weeks of evolution; **(B)** NMDS ordination plot depicting the pair-wise **Bray-Curtis dissimilarity between soil microcosms after six weeks of incubation (stress** = 0.096). Points represent individual microcosms belonging to the control (open circles) and copper (black circles) treatment, such that microcosms similar in their genus-level composition are ordinated closer together; **(C)** Relative abundance of the ten most common genera and their mean siderophore production. Genera are listed in order of their mean across-treatment siderophore production, increasing from top to bottom, such that blue- and red genera are non-producers and producers, respectively; **(D)** The effect of copper background (filled and open symbols are presence and absence of copper contamination, respectively) on metal tolerance, where more negative values indicate astronger inhibitory effect of 6.17 mM CuSO_4_ on bacterial growth. Bars denote 1SE.

We identified clones at the genus-level to explore the role of species sorting in driving siderophore production. Community composition varied significantly between treatments (PERMANOVA: *F*_*1, 11*_ = 3.88, *P* = 0.015; multivariate dispersion similar across treatments: *F*_*1,11*_ = 0.021, *P* = 0.91; Fig. 4B), with siderophore-producing genera being selectively favoured in copper-contaminated compost (Fig. 4C). Crucially, clones isolated from the copper treatment were significantly less inhibited when grown in toxic copper broth compared to those from the control treatment (LME: χ^2^ = 6.80; d.f. = 1; *P* < 0.01; Fig. 4D), which was mediated by increased siderophore production (LME: χ^2^ = 16.68; d.f. = 1; *P* < 0.001).

## DISCUSSION

In this study, we investigated how heavy metals affected ecological selection for siderophore production – an interspecific microbial public good – across an environmental gradient of contaminated soil and during a controlled experiment in compost. We hypothesised there could be selection for both increased and decreased siderophore production, because of the detoxifying effect of siderophores and the potential for interspecific siderophore exploitation, respectively. Our findings suggest that the presence of toxic metals resulted in ecological selection for taxa that produced large amounts of siderophore. We also confirmed that bacteria investing more in siderophores suffered less growth inhibition in the presence of toxic copper.

Ecological selection for increased siderophore production contrasts with previous in vitro within-species (*P. aeruginosa*) results, in which non-producing ‘cheats’ were able to outcompete siderophore producers in copper-contaminated broth (O’Brien *et al.* 2014). A key reason for this difference is likely to be the spatial structure in soil/compost resulting in localised detoxification, such that producers and their immediate neighbours gain the most from their siderophores (Hamilton 1964; West & Buckling 2003; Buckling *et al.* 2007; West *et al.* 2007; Lujan *et al.* 2015). Hence, low siderophore producers should experience more of the toxic metal effect. Limited dispersal would also lead to immediate neighbours having a higher probability of being conspecifics - a likely reason as to why taxa that typically produce more siderophores dominated the community when exposed to toxic metals. Direct comparison of intra- and inter-specific changes in siderophore production in soil would tease apart the differing roles of spatial and community structure in determining these results.

In our survey of a former mining area, soil acidity and total metal contamination positively co-varied, with both prolonged metal leaching in acidic soils and precipitation in more basic soils likely contributing to this pattern (Alloway 1990; Adriano 2001). This covariance may well have contributed to the patterns we observed. First, acidity is a major determinant of microbial diversity and composition (e.g., Fierer & Jackson 2006; Griffiths *et al.* 2011), hence pH-mediated selection may have indirectly favoured taxa that produce siderophores in larger amounts. Second, acidity affects metal speciation and bio-availability to microbes in variable ways (Lofts *et al.* 2004; Gobran & Huang 2011), with iron becoming largely insoluble at pH > 6.5 (Guerinot 1994). As such, increased siderophore production in basic soils, which also had the highest metal concentrations, may have been driven by selection imposed by iron limitation. However, our experimental manipulations, where the same compost community was propagated in both the presence and absence of copper, strongly suggest a direct effect of heavy-metal imposed selection on siderophore production. This manipulation did have a small effect on pH (copper decreased pH from approximately 7.1 to 6.6), but in this case there was negative, rather than positive, covariance between pH and metal contamination.

It was initially surprising to find that microbial densities were similar along the natural contamination gradient; several studies have demonstrated that toxic heavy metals reduce microbial abundance (reviewed in Giller *et al.* 1998). These differences may reflect the relatively short time scale of exposure in most studies as well as low metal concentrations. Given the mining history of the site used in this study, microbes are likely to be relatively well adapted to the toxic conditions, through other more direct resistance mechanisms not investigated here (Nies 1999; Bruins *et al.* 2000; Valls & De Lorenzo 2002), in addition to siderophore production. As a consequence of this resistance, microbial communities were perhaps able to reach high densities in the presence of normally toxic metal concentrations. It is important to emphasise that selection of taxa with increased copper tolerance occurred very rapidly in our experiment, although as stated above we can’t rule out a role of additional resistance mechanisms that positively co-vary with siderophore production.

Human-imposed heavy metal contamination is a major problem for natural ecosystems and several studies have noted that addition of siderophores or siderophore-producing microbes could aid in detoxifying contaminated soils, particularly when combined with the use of heavy metal hyper-accumulating plants (Lebeau *et al.* 2008; Dimkpa *et al.* 2009). Our results provide some key insights into the optimal use of siderophores for remediation. The addition of high siderophore-producing bacteria following recent contamination events is likely to be effective, because these organisms should have a selective advantage and hence contribute to increasing the community-level production of siderophores. However, their addition is unlikely to significantly improve remediation of historically contaminated sites, in which siderophore production will already have been stabilised by selection. The direct addition of siderophores, while providing a short-term benefit, may actually result in a longer-term negative effect on remediation regardless of length of time since contamination, as selection for siderophore production is relaxed. More generally, our results highlight that interspecific public goods production can be maintained at high levels in natural microbial communities, despite the potential of exploitation by cheating non-producers.

## ACKNOWLEDGEMENTS

We like to thank Pawel Sierocinski, Uli Klumper and Chris Bryan for discussions on bioinformatics and metals. This work was funded by the AXA Research Fund, BBSRC and NERC to AB. SOB was funded by a “Bridging the Gaps” award and PhD scholarship from the University of Exeter. NT was funded by the European Union’s Horizon 2020 research and innovation programme under the Marie Sklodowska-Curie grant agreement No 656647. AML was supported by Marie Curie International Incoming Fellowships within the European Union Seventh Framework Programme. AB acknowledges support from the Royal Society.

Supplementary information is available at the Ecology Letters journal website.

